# Mucosal vaccination with cyclic-di-nucleotide adjuvants induces effective T cell homing and IL-17 dependent protection against *M. tuberculosis* infection

**DOI:** 10.1101/2020.11.25.398651

**Authors:** Robyn M. Jong, Erik Van Dis, Xammy Nguyenla, Alexander Baltodano, Gabrielle Pastenkos, Chenling Xu, Nir Yosef, Sarah M. McWhirter, Sarah A. Stanley

**Affiliations:** Molecular and Cell Biology, Division of Immunology and Pathogenesis, University of California, Berkeley, Berkeley, California, United States of America; School of Public Health, Division of Infectious Disease and Vaccinology, University of California, Berkeley, Berkeley, California, United States of America; Aduro Biotech, Inc., 740 Heinz Avenue, Berkeley, California, United States of America; Comparative Pathology Laboratory, University of California, Davis, Davis, California, United States of America; Center for Computational Biology, University of California, Berkeley, Berkeley, California, United States of America; Department of Electrical Engineering and Computer Sciences, University of California, Berkeley, Berkeley, California, United States of America; Ragon Institute of MGH, MIT and Harvard, Cambridge, MA, United States of America; Chan-Zuckerberg Biohub Investigator

## Abstract

The only licensed vaccine for tuberculosis, *Mycobacterium bovis* Bacille Calmette-Guérin (BCG), is not reliably effective against adult pulmonary tuberculosis. A major hurdle to tuberculosis vaccine development is incomplete understanding of successful immunity against the causative agent *Mycobacterium tuberculosis*. Recently, we demonstrated that a protein subunit vaccine adjuvanted with STING-activating cyclic-di-nucleotides (CDNs) robustly protects against tuberculosis infection in mice. Here we show mucosal vaccination with this vaccine induces production of T cells that home to lung parenchyma and penetrate lesions in the lung. Protection is partially dependent on IL-17, type I interferon (IFN), and IFN-γ, while the transcription factor STAT-6 is dispensable. Single cell transcriptomics reveals mucosal vaccination with a CDN vaccine increases transcriptional heterogeneity in CD4 cells, including a significant population of non-classical IFN-γ and IL-17 co-expressing Th1-Th17 cells, as well as markers of memory and activation. Th1-Th17 cells in vaccinated mice are enriched for expression of the T cell functional markers *Tnfsf8* and *Il1r1* relative to more conventional Th1 cells. These data provide critical insight into the immune mediators and diversity of T cell responses that can contribute to vaccine efficacy against *M. tuberculosis* infection.

## Introduction

Tuberculosis (TB) causes more deaths worldwide than any other single infectious agent^1^. The existing live-attenuated vaccine against TB, *M. bovis* Bacille Calmette-Guérin (BCG), is widely administrated in many countries with endemic TB and is effective against severe forms of childhood disease^1^. However, efficacy wanes over time and BCG provides minimal protection against adult pulmonary TB^2^. There is a clear need for an improved vaccine to combat this global health threat. One of the barriers to the development of effective vaccines for TB is the lack of reliable immune correlates of protection. Immunity to TB has long been considered to be dependent on the production of IFN-γ by CD4 T cells^3^, as defects in IFN-γ signaling are linked to susceptibility to mycobacterial infection^4^. Thus, the development of new TB vaccines to date has focused on eliciting IFN-γ-producing CD4 T cells and polyfunctional CD4 T cells that produce IFN-γ, TNF-α, and IL-2^5^. A highly anticipated clinical trial of the BCG booster protein subunit vaccine MVA-85A showed successful induction of *M. tuberculosis*-specific polyfunctional CD4 T cells in infants^6^. However, this immune response did not translate into significant protection against TB infection or disease^6^. More recently, the protein subunit vaccine candidate M72/AS01E stimulated IFN-γ producing T cells and demonstrated 50% efficacy in preventing the transition from latent to active disease over a 3-year period^7^. Why two vaccines that both elicit IFN-γ-producing CD4 T cells bore different results is unclear, however these results suggest that alternate biomarkers of efficacy may more clearly predict vaccine efficacy.

Animal studies have suggested the existence of IFN-γ-independent mechanisms of CD4 T cell-mediated control of *M. tuberculosis* infection, however, the basis of this control has remained elusive^8^. IL-17 producing Th17 T cells do not contribute to control of primary TB infection in mice, possibly because they are not robustly elicited during primary infection. However, intranasal administration of BCG^9,10^ or protein subunit vaccines^11,12^ elicits IL-17-producing memory T cells that correlate with enhanced protection against *M. tuberculosis* challenge in rodent models. Th17 cells are a heterogenous effector subset of CD4 T cells that can range in function from inflammatory to regulatory and can even transdifferentiate into Th1 cells^13^. Although TB vaccine-elicited memory Th17s can persist in lung-draining lymph nodes and acquire Th1 characteristics after challenge^14^, it is unclear which Th17 subtype is most important for vaccine-elicited control of TB, or how this subtype enhances control of *M. tuberculosis* infection.

In order to exert control of infection, T cells must both differentiate into a protective effector state and adopt an effector phenotype that allows them to migrate to the site of infected macrophages in the lung. In mouse models, lung parenchyma-homing CXCR3^+^ CD153^+^ KLRG1^-^ CD4 T cells mediate superior protection against *M. tuberculosis* compared to vasculature-associated KLRG1^+^ CD4 T cells that do not migrate into the lung tissue^15,16^. Furthermore, T cells that are blocked from entering the granuloma core are unable to provide as effective protection as T cells that penetrate into the central core of infected macrophages^17^. However, it is unclear how to rationally tailor a vaccine that will elicit CD4 T cell homing to granuloma cores.

We recently demonstrated in mice that an experimental protein subunit vaccine formulated with STING-activating cyclic-di-nucleotide (CDN) adjuvants is highly efficacious in mice, particularly when delivered by a mucosal route^18^. Here we find that protective efficacy correlates with the production of parenchymal homing T cells, penetration of T cells into lung lesions, and a T cell response dominated by Th17 at early timepoints. We further show that type I IFN, IL-17, and IFN-γ are required for complete intranasal CDN-adjuvanted vaccine-induced protection. While intranasal vaccine delivery elicits IFN-γ-producing CD4 T cells, vaccine-mediated protection is partially IFN-γ-independent and does not correlate with increased Th1 responses. Single cell transcriptomics (scRNA-seq) shows that naïve infected mice predominantly produce KLRG1^+^ CX3CR1^+^ effector Th1s that do not infiltrate lung lesions, while mucosal vaccination promotes activation and memory marker-expressing CD4 T cells. Intranasal vaccination induces T cell heterogeneity, particularly within the *Il17a*-expressing subset, and a large number of mixed Th1-Th17 cells. These results demonstrate that CDN-adjuvanted protein subunit vaccines generate a multifaceted CD4 T cell response that leads to enhanced protection against TB disease.

## Results

We previously found that mucosal administration of a protein subunit vaccine with CDN adjuvants elicits significant protection against *M. tuberculosis* in the mouse model of infection^18^. We therefore set out to define the mechanisms underlying the efficacy of CDN adjuvants for TB vaccines. For antigen we used either 5Ag—a five antigen fusion of *M. tuberculosis* proteins that includes the immunodominant proteins Ag85B and ESAT-6 and three proteins predicted to be expressed in bacterial latency^18^—or H1 antigen, a fusion of Ag85b-ESAT6 alone, as 5Ag and H1 were functionally equivalent for protection out to 4 weeks post challenge^18^ (Fig. S1). Protein antigen was formulated with the CDN ML-RR-cGAMP^19^.

### CDN-adjuvanted protein subunit vaccines elicit a rapid influx of T cells and control of infection upon *M. tuberculosis* challenge

We first characterized the kinetics of protection and CD4 T cell infiltration elicited by H1/ML-RR-cGAMP in lungs after challenge. Mice were vaccinated 12 weeks prior to challenge with *M. tuberculosis*, either by the intranasal route (i.n.) with H1/ML-RR-cGAMP three times at 4 week intervals, or once with subcutaneous (s.c.) BCG, the standard route for BCG. Vaccination with H1/ML-RR-cGAMP resulted in increased protection when compared with BCG at 1 and at 2 weeks post challenge, demonstrating that mucosal CDN adjuvanted-vaccination induces immunity earlier than BCG vaccination (Fig. 1a). Protection correlated with an influx of antigen-specific IL-17-producing T cells into the lungs at 1 week post challenge that increased steadily until 3 weeks post challenge (Fig. 1b). Neither mock (PBS-treated) nor BCG-vaccinated mice showed appreciable Th17 responses (Fig. 1b). Th1 frequency dramatically increased in mock vaccinated mice between 2-3 weeks post challenge (Fig. 1c), eclipsing that of the vaccinated mice and coinciding with a larger bacterial burden (Fig. 1a). A smaller population of CD4 T cells produced both IFN-γ and IL-17 (Fig. 1d). While several sources have suggested that TCRγ/δ T cells are the main source of IL-17 in *M. tuberculosis*-infected mouse lungs during primary infection^20,21^, we observed no difference in frequencies of TCRγ/δ^+^ T cells or in the subset of IL-17^+^ TCRγ/δ T cells in mock vs vaccinated mice after infection (Fig. S2a, b). Thus, the evidence points to a role for vaccine-induced IL-17-producing conventional CD4 T cells in early and sustained mucosal vaccine efficacy.

**Figure 1.**
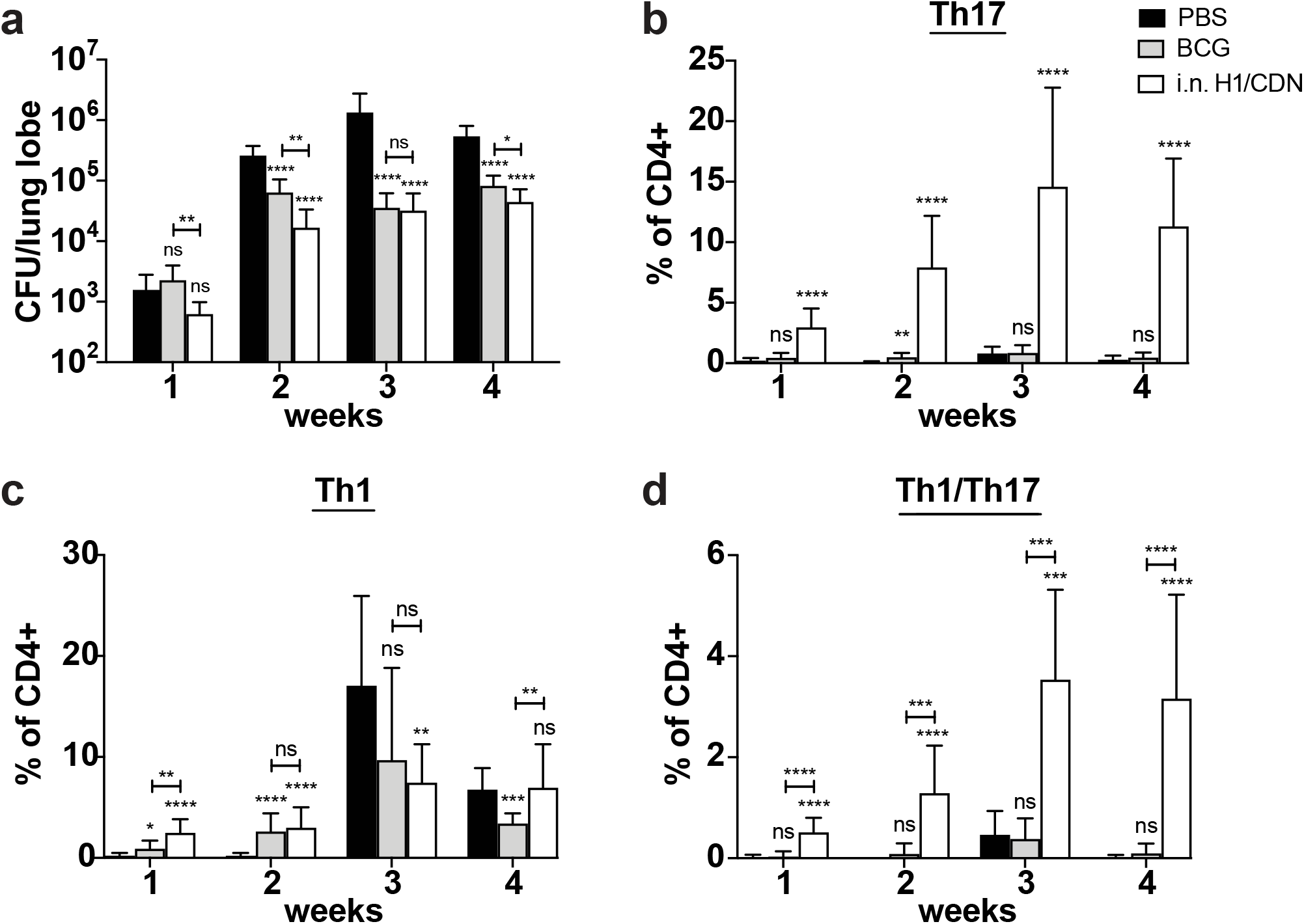
Intranasal immunization induces a Th17 response that precedes significant Th1 influx into the lungs of *M. tuberculosis*-challenged mice. Mice were vaccinated once with subcutaneous *M. bovis* BCG 12 weeks before challenge or primed with intranasal *M. tuberculosis* H1 antigen and ML-RR-cGAMP cyclic-di-nucleotide adjuvant (i.n. H1/CDN) 12 weeks before challenge, then boosted twice at 4 week intervals after priming, before *M. tuberculosis* aerosol challenge with 50-100 CFU. Mice were mock primed and boosted with PBS as a control. (**a**) Lung bacterial burden was enumerated at 1 – 4 weeks post *M. tuberculosis* challenge. (**b-d**) Intracellular cytokine staining (ICS) for percentage of lung CD4 T cells that produce (**b**) IL-17 (Th17), (**c**) IFN-γ (Th1), or (**d**) both IFN-γ and IL-17 (Th1/Th17) after restimulation *ex vivo* with recombinant Ag85b peptide pool. Data are expressed as mean (± SD) of eight to ten mice per group from two independent experiments. Mann-Whitney t test p values calculated in comparison to PBS vaccinated controls except where indicated; *p ≤ 0.05, **p ≤ 0.01, ***p ≤ 0.001, ****p ≤ 0.0001. Significance is relative to PBS control unless otherwise indicated.

### IL-17 is required for full efficacy of CDN adjuvanted protein subunit vaccines

To determine whether IL-17 is necessary for protective efficacy, we vaccinated wild type and *Il17*^−/−^ mice and enumerated bacterial burdens in the lungs 4 weeks post challenge with *M. tuberculosis*. Bacterial burden was unaffected by the loss of IL-17 in mice that were mock vaccinated, vaccinated s.c. with 5Ag/ML-RR-cGAMP, or vaccinated s.c. with BCG (Fig. 2a). However, we observed a significant loss of protection in *Il17*^−/−^ mice that were vaccinated with i.n. 5Ag/ML-RR-cGAMP. While IL-17 has been reported to impact the development and recruitment of vaccine-induced Th1 cells^22,23^, Th1 cell production was unaffected in *Il17*^−/−^ mice at this timepoint (Fig. 2b). These data suggest that the efficacy of mucosal CDN vaccines is partially IL-17 dependent.

**Figure 2.**
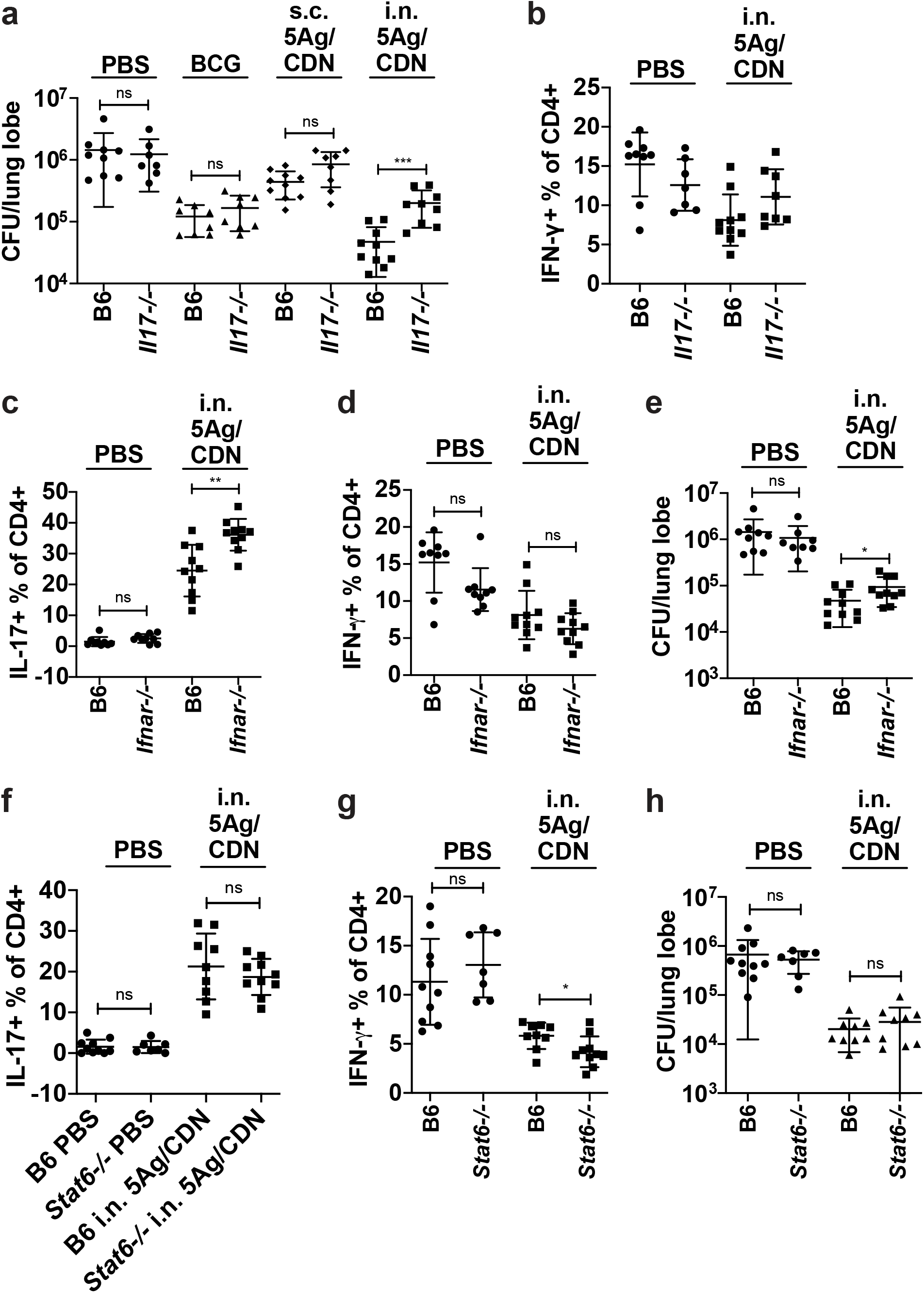
IL-17 and Type I IFN, but not STAT6, are essential for full mucosal vaccine efficacy. Wild-type C57BL/6 (B6) and *Il17*^*–/–*^ mice were vaccinated once with subcutaneous *M. bovis* BCG 12 weeks before *M. tuberculosis* aerosol challenge, or primed with either subcutaneous or intranasal *M. tuberculosis* 5Ag antigen and ML-RR-cGAMP cyclic-di-nucleotide adjuvant (i.n. 5Ag/CDN) 12 weeks before challenge, followed by two boosts. (**a, e, h**) At 4 weeks post challenge, mouse lungs were harvested and analyzed for bacterial burden. **(b-d, f, g)** ICS analysis of *ex vivo* restimulated lung leukocytes for **(c, f)** % IL-17^+^ and **(b, d, g)** % IFN-γ^+^ CD4 T cells. Data are expressed as mean (± SD) of eight to ten mice per group from two independent experiments. Mann-Whitney t test p values; *p ≤ 0.05, **p ≤ 0.01, ***p ≤ 0.001.

### Type I interferon and IFN-γ contribute to CDN adjuvanted mucosal vaccine efficacy

STING agonists strongly induce type I IFN expression through IRF3-dependent signaling^24^. As type I IFN responses can negatively regulate the development of Th17 cells^25^ and type I IFN can play a detrimental role in primary *M. tuberculosis* infection immunity^26,27^, it is somewhat surprising that STING agonists elicit robust Th17-dependent protective immunity. We tested the efficacy of i.n. 5Ag/ML-RR-cGAMP vaccination in IFNα/β receptor deficient mice (*Ifnar*^*–/–*^). As expected, the Th17 response was more robust in the lungs of *Ifnar*^*–/–*^ mice at 4 weeks post challenge (Fig. 2c), while the Th1 response was unaffected (Fig. 2d). However, despite an increased *M. tuberculosis*-specific Th17 response, we observed a small but statistically significant loss of control in vaccinated *Ifnar*^*–/–*^ mice (Fig. 2e). The increased Th17 response in *Ifnar*^*–/–*^ mice thus cannot compensate for the loss of type I IFN in vaccine-elicited protective immunity.

STING also signals through both NF-κB and STAT6. STAT6 is a transcription factor that can be activated downstream of STING, leading to chemokine production that promotes control of viral infections^28^. Loss of STAT6 did not affect Th17 responses in *Stat6*^*–/–*^ vaccinated mice (Fig. 2f). A slight decrease in Th1 frequency in *Stat6*^*–/–*^ intranasal vaccinated mice (Fig. 2g) did not translate to an impact on the efficacy of i.n. 5Ag/ML-RR-cGAMP (Fig. 2h). Therefore, the efficacy of CDN adjuvanted TB vaccines is not dependent on STAT6 signaling.

We next sought to determine whether the classical Th1 response plays a significant role in protection. We vaccinated mice lacking IFN-γ (*Ifng*^−/−^) with H1/ML-RR-cGAMP subcutaneously or intranasally and harvested lungs at 3 weeks post challenge to assess vaccine efficacy. Loss of IFN-γ did not affect the robust Th17 response in i.n. vaccinated mice prior to challenge (Fig. S3) or at the peak of the T cell response at 3 weeks post challenge (Fig. 3a). While all *Ifng*^−/−^ mice were more susceptible to *M. tuberculosis* infection, the subcutaneous and mucosal vaccines both provided significant protection compared to mock vaccination (Fig. 3b, c). While this demonstrates that both routes of inoculation elicit IFN-γ-independent mechanisms of control, the magnitude of protection was significantly reduced in *Ifng*^−/−^ mice (Fig. 3c), indicating that IFN-γ does play some role in vaccine efficacy, in conjunction with IFN-γ-independent mechanisms.

**Figure 3.**
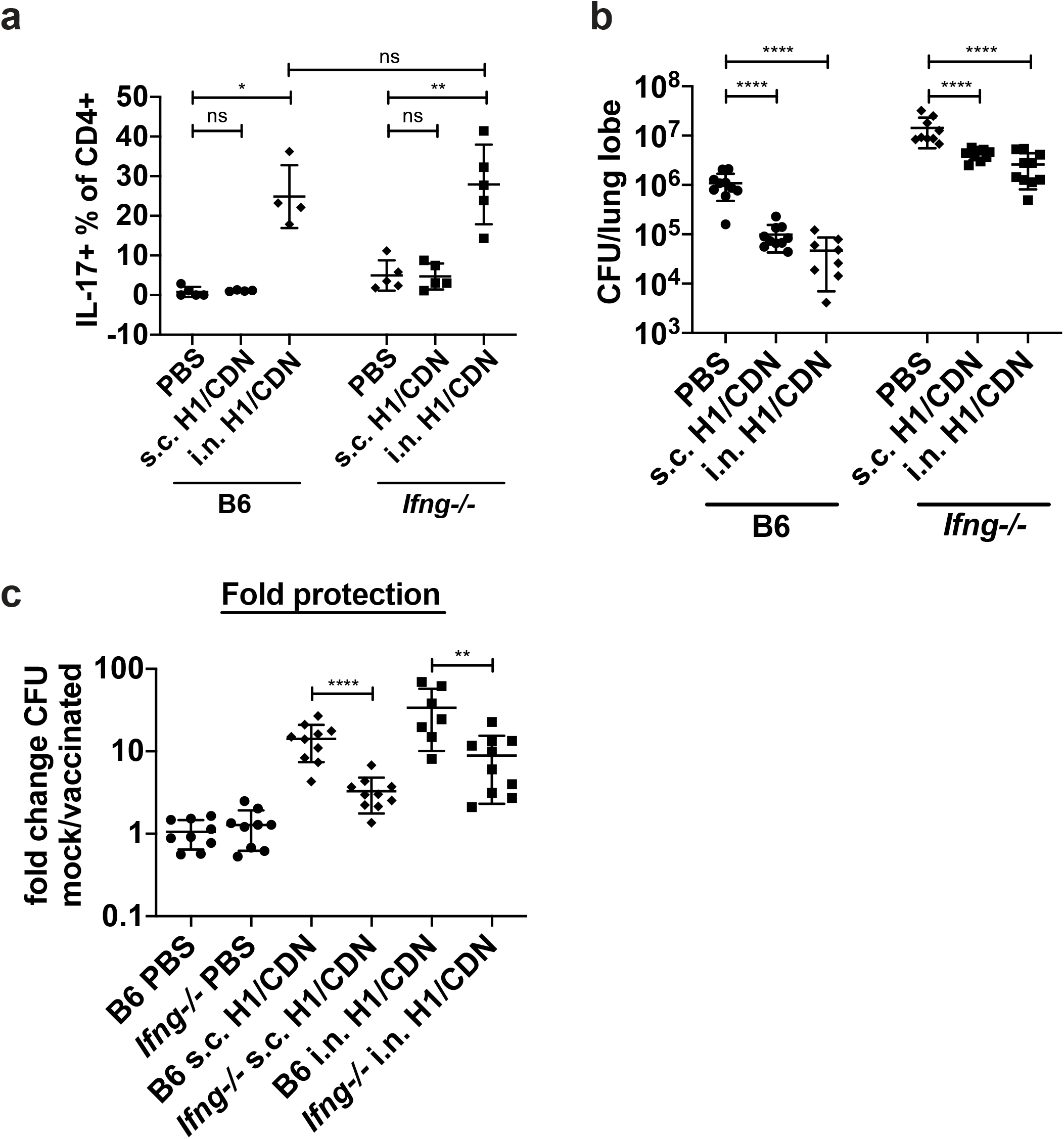
Vaccine efficacy is not fully dependent on the presence of IFN-γ. **(a)** ICS analysis of *ex vivo* restimulated lung leukocytes from mock or i.n. H1/ML-RR-cGAMP (i.n. H1/CDN) vaccinated B6 or *Ifng*^*–/–*^ mice 4 weeks post *M. tuberculosis* challenge for % IL-17^+^ CD4 T cells. **(b)** CFU counts from lungs of vaccinated mice 4 weeks post challenge. **(c)** Fold protection of vaccination, calculated as the fold-decrease in lung bacterial burden between vaccinated and mock vaccinated mice. Data are expressed as mean (± SD) of five to ten mice per group from (**a**) one representative experiment or (**b, c**) two independent experiments. Mann-Whitney t test p values; *p ≤ 0.05, **p ≤ 0.01.

### Lymphocytes home to infected tissue in CDN vaccinated mice

Proper localization of T cells in infected lungs is crucial for effective T cell-mediated immunity^29^. CXCR3^hi^ CD4 T cells can localize to the infected mouse lung parenchyma and mediate superior bacterial resistance compared to KLRG1^hi^ T cells, which remain trapped in the vasculature^16,30^. We found that i.n. vaccination with H1/ML-RR-cGAMP induced higher frequencies of CXCR3^+^ and lower frequencies of KLRG1^+^ CD4 T cells in infected mouse lungs than s.c. BCG vaccination or mock vaccination (Fig. 4a,b). Conversely, BCG vaccination induced a higher percentage of KLRG1^+^ cells than i.n. H1/ML-RR-cGAMP vaccination (Fig. 4b). In naïve infected mice, significant numbers of KLRG1^+^ T cells appeared in the lungs at 3-4 weeks post *M. tuberculosis* challenge (Fig. 4b). We next sought to determine whether i.n. vaccination with 5Ag/ML-RR-cGAMP changed T and B cell localization within infected lung tissue. Consistent with previous reports^17^, we found that in naïve infected mice, very few CD3^+^ T cells were observed in lung lesions (Fig. 4c). However, i.n. vaccination effectively enhanced infiltration of CD3^+^ T cells into the lesion (Fig. 4c, d) without increasing overall lung inflammation (Fig. S4a). Although previous reports associated the efficacy of mucosal TB protein subunit vaccines with the formation of B cell follicles in vaccinated murine lungs^11,12^, we did not observe B cell follicles with germinal center organization in any of the mice at 4 weeks post challenge (Fig. S4b). However, i.n. vaccinated mouse lungs were more likely to contain lymphocyte aggregates without germinal centers (lymphoid nodules, Fig. S4c) and B220^+^ staining was increased in i.n. vaccinated mouse lungs (Fig. 4e, f), suggesting that mucosal vaccination may boost the B cell response. Overall levels of inflammation did not correlate with protection (Fig. S4a-g). Thus, mucosal vaccination leads to intralesional T homing and B cell infiltration into infected lungs.

**Figure 4.**
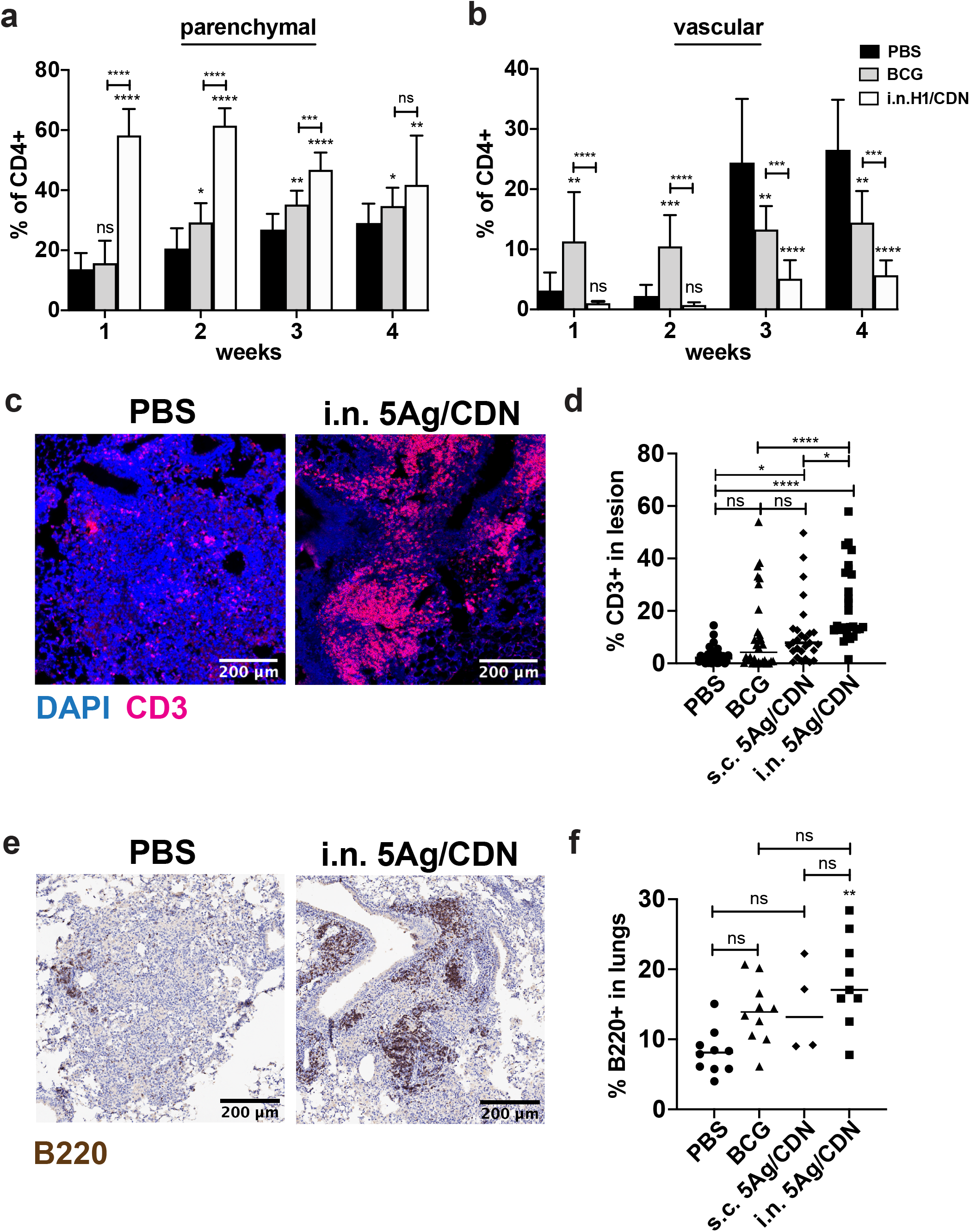
Lymphocytes home to lung lesions in mucosal vaccinated mice. **(a)** Surface staining for percentage of CXCR3^+^ KLRG1^-^ lung CD4 T cells from mock and i.n. H1/ML-RR-cGAMP (i.n. H1/CDN) vaccinated mice. **(b)** Surface staining for percentage of CXCR3^-^ KLRG1^+^ lung CD4 T cells. **(c)** Representative immunofluorescent staining of formalin-fixed, paraffin-embedded lung sections from mock and 5Ag/ML-RR-cGAMP vaccinated mice for T cell marker CD3 and nuclear DAPI stain. **(d)** Quantification of CD3^+^ immune cells out of total cells in lung lesions. **(e)** Representative immunohistochemical staining of lung sections for B cell marker B220. **(f)** Quantification of B220^+^ lung area. Data are expressed as mean (± SD) of eight to ten mice per group from two independent experiments. (**a, b**) Mann-Whitney t test p values; (**d, f**) Kruskal Wallis test followed by Dunn multiple comparison posthoc p-values; *p ≤ 0.05, **p ≤ 0.01, ***p ≤ 0.001, ****p ≤ 0.0001. Significance is relative to PBS control unless otherwise indicated.

### Single cell analysis reveals shifts in naïve and vaccinated CD4 T cell compartment transcriptomes

To gain a more detailed understanding of the characteristics of the CD4 T cell response during *M. tuberculosis* infection we performed single cell RNA-sequencing (scRNA-seq)^31^ on CD4 T cells sorted from naïve and i.n. H1/ML-RR-cGAMP vaccinated infected mouse lungs at the peak of the T cell response, 3 weeks post infection. The filtered dataset, analyzed by single cell variational inference (scVI)^32^, consisted almost exclusively of CD4 T cells (Fig. S5a). We first characterized the lung CD4 T cell response in naïve infected animals. Using signature transcription factor, cytokine, and cell surface marker expression, we were able to assign 56% of cells to classic CD4 T cell subsets while the rest of the cells could not be confidently assigned due to lack of marker expression. Our analysis identified clear populations of Th1 (*Ifng*), Treg (*Foxp3*), and naive (*Ccr7, Sell*; *Cd44*^*lo*^) T cells (Fig. 5a). The largest population of assigned cells expressed the Th1 gene marker *Ifng*, in agreement with our flow cytometry analysis (Fig. 1c). Very few T cells expressed the Th17 marker *Il17a*, and a small population of cells expressed *Foxp3* (Fig. 5a, c). Few Th2 (*Gata3, Il4*) or Tfh (*Bcl6, Cxcr5*) cells were observed. Naive, Treg, and proliferating (*Mki67*) cells formed a clear cluster, while other cell subtypes—Th2, Th17, Tfh—varied more continuously in the transcriptional space. CD4 T cells were also isolated from i.n. H1/ML-RR-cGAMP vaccinated infected lungs, and 46% of cells were assigned to a CD4 T cell subset (Fig. 5b). *Ifng*-expressing Th1 cells were the dominant CD4 subtype in vaccinated lungs, although fewer in frequency compared to the mock vaccinated sample (Fig. 5a, b). As expected (Fig. 1c, d), a significant population of CD4 T cells in mucosally vaccinated mice expressed *Il17a*. 20-30% of *Il17a*-expressing cells in vaccinated mice also expressed *Ifng* (Fig. 5b), suggesting that these cells—denoted here as Th1-Th17—were not classical Th17 cells. Th17 signature transcription factor *Rorc* expression, but not Th1 signature transcription factor *Tbx21* expression, was detected in Th1-Th17 cells (Fig. S5b). Thus, the CD4 T cell compartment in vaccinated infected mice is characterized by a new population of *Il17a*-expressing cells that is not apparent in naïve infected mice, and a portion of these Th17 cells co-express *Ifng*.

**Figure 5.**
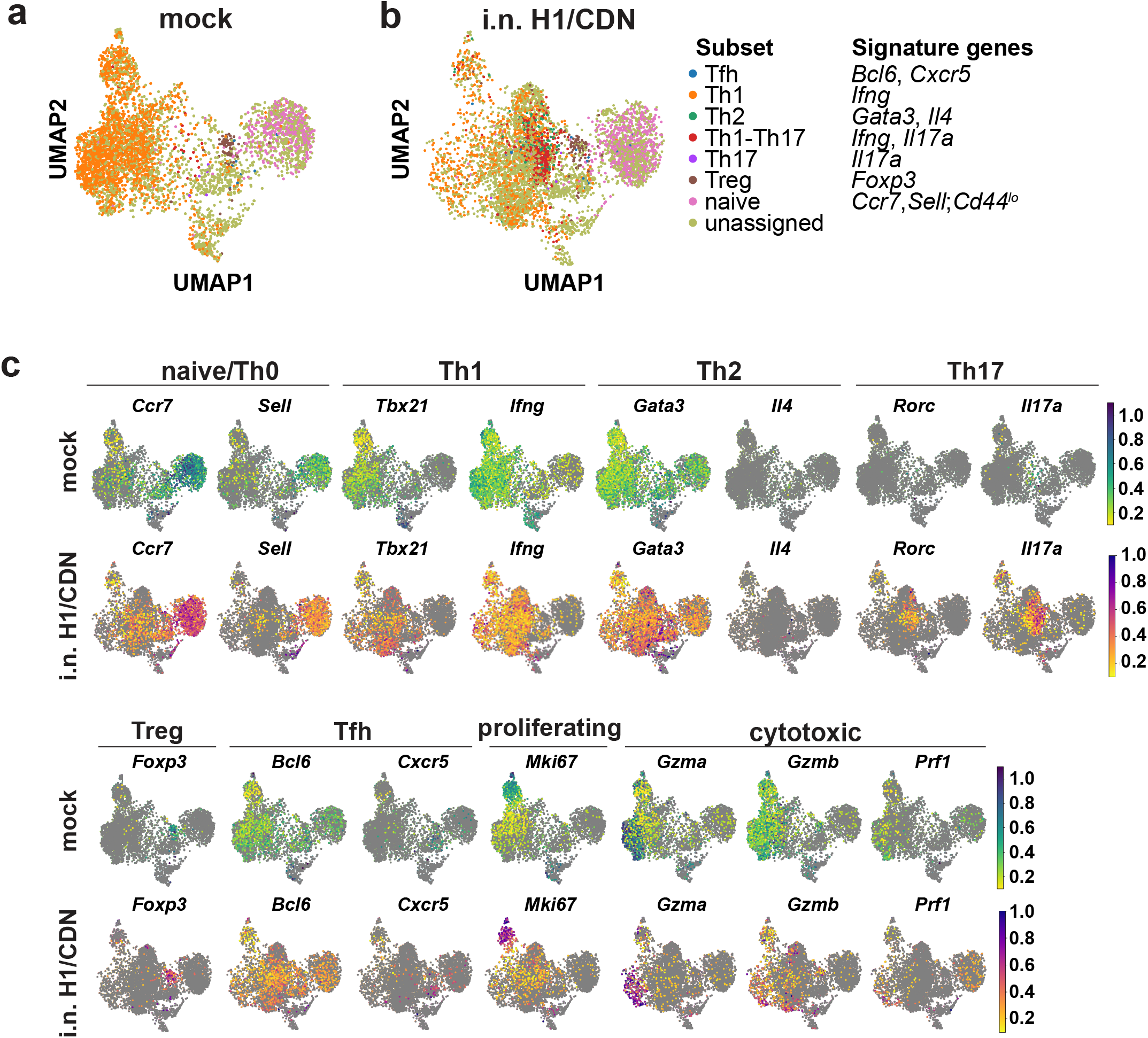
Single cell transcriptional analysis reveals the effects of mucosal vaccination on mouse lung CD4 T cell compartments. Subsets of CD4 T cells were manually annotated based on signature expression of lineage-specific transcription factor, cytokine, and cell surface markers. **(a)** UMAP plots of CD4 T cell subsets isolated from naïve (mock) or **(b)** i.n. H1/ML-RR-cGAMP (i.n. H1/CDN) vaccinated mice 3 weeks post infection. **(c)** Signature gene expression of CD4 T cell subtypes in naïve (mock) and i.n. H1/ML-RR-cGAMP vaccinated mice. The color scale indicate the percentile expression level for each cell that has non-zero expression.

Signature gene expression was generally similar between mock and mucosally vaccinated CD4 T cell subtypes (Fig. S5b). To determine whether functionality differed between the CD4 T cell compartments, we first identified genes upregulated in unvaccinated mice. *Ifng, Klrg1*, and *Cx3cr1* expression was increased in CD4 T cells from naïve infected lungs compared to vaccinated infected lungs (Fig. 6b and Fig. S5b), as expected (Fig. 1c and Fig 4b, c). Interestingly, *Nkg7*, a regulator of cytotoxic granule exocytosis, was highly expressed in the bulk CD4 population of naïve infected lungs (Fig. 6a, b), as were cytotoxic markers *Gzma, Gzmb*, but not *Prf1* (Fig. 6b). These expression patterns were also evident in the *Ifng*-expressing subsets in both mock and mucosally vaccinated infected mice (Fig. 6c). Mucosal vaccination led to upregulation of genes associated with the *Rorc*-expressing Th17 lineage (Fig. 6d), such as *Tmem176a, Tmem176b*, and *JunB*. The naïve and central memory T cell marker *Il7r* was highly expressed in the i.n. vaccinated CD4 T cell compartment (Fig. 6d), but not naïve T cell markers *Ccr7* and *Sell* (Fig. S5c, d). Increased *Rbpj* and *Cd44* expression in vaccinated mice (Fig. 6d) may contribute to survival of effector and memory T cells and memory Th1 development. Other activation marker genes such as *Il2ra, Cd69, Ctla4* (Fig. 6d, e), but not *Pdcd1* (Fig. S5c, d), also had increased expression within the bulk CD4 population and *Ifng*-expressing Th1 subset. Vaccination did not affect proliferation marker *Mki67* expression (Fig. S5c, d). Within the Th1 subset, the top 10 differentially expressed genes identified—including *Klrg1, Cx3cr1, Nkg7, Il7r, Tmem176a, Tmem176b*—were also identified as differentially expressed in the bulk CD4 population (Fig. S5e), suggesting that the bulk CD4 population differential expression is driven by transcriptional changes in the Th1 subset rather than changes in abundance of T cell subsets. The magnitude of the different expression patterns was either similar to the bulk CD4 population, or was amplified in the Th1 subset (Fig. 6d, e and Fig. S5d). Thus, single cell transcriptomics reveals that primary challenge with *M. tuberculosis* in unvaccinated mice induces lung CD4s that transcriptionally resemble terminally differentiated effector KLRG1^+^ CX3CR1^+^ Th1 cells that contribute little to control of *M. tuberculosis*. In contrast, mucosal vaccination elicits a more diverse CD4 T cell compartment with Th17, activation, and memory T cell markers.

**Figure 6.**
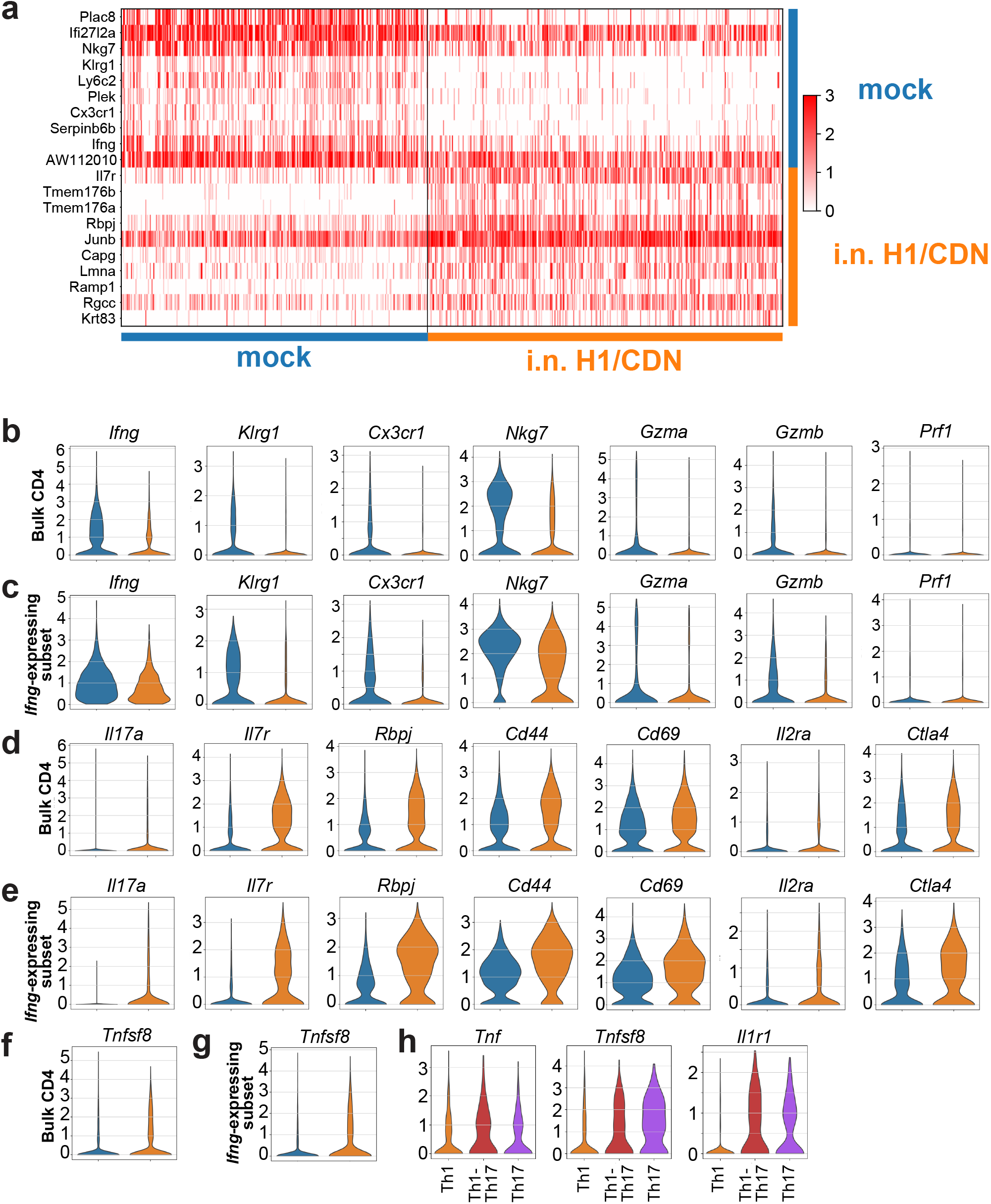
CD4 T cell transcriptomes from naïve infected mice show increased terminally differentiated Th1 marker expression, while mucosal vaccination leads to memory marker expression and the development of Th1-Th17 and Th17 cells. Top differentially expressed genes in naïve (mock) and i.n. H1/ML-RR-cGAMP (i.n. H1/CDN) vaccinated mouse CD4 T cells obtained by Student’s t-test with overestimated variance. **(b, c)** Markers of Th1 lineage, terminal differentiation, and cytotoxicity in **(b)** bulk and **(c)** *Ifng*-expressing CD4 populations in naïve (blue) and i.n. H1/ML-RR-cGAMP vaccinated (orange) infected mouse lungs. **(d, e)** Markers of Th17 lineage, memory, and activation in **(d)** bulk and **(e)** *Ifng*-expressing CD4 populations. **(f, g)** *Tnfsf8* expression in **(f)** bulk and **(g)** *Ifng*-expressing CD4 populations. **(h)** Markers of polyfunctionality, protective capacity, and effector function in Th1, Th1-Th17, and Th17 subsets.

Polyfunctional T cells expressing IFN-γ, TNF, and IL-2 have been implicated in vaccine-induced protection against TB^5^. Neither *Ifng* (Fig. 6b) nor *Il2* (Fig. S5c, d) expression was significantly increased with mucosal vaccination, but *Tnf* expression was slightly elevated in the mucosal vaccinated T cell compartment (Fig. S5c, d), particularly within the Th1-Th17 and Th17 subsets (Fig. 6h). Besides *Ifng* expression, Th1-Th17 transcriptomes seemed much closer to transcriptomes from Th17s expressing *Il17a* alone than those of classical Th1s (Fig. S5f). Intriguingly, mucosal vaccine-induced CD4 T cells expressed higher levels of the gene encoding CD153, *Tnfsf8* (Fig. 6f, g), a marker for protective lung-homing T cells in mice^15^ and humans^33^. Cells expressing *Il17a* but not *Ifng* expressed the highest levels of *Tnfsf8*, as well as *Il1r1*, a critical component of effector T cell function (Fig. 6h). CD153 and IL-1R1 may therefore promote the development or function of protective CD4 T cells in mice vaccinated with CDN-adjuvanted vaccines.

Many cells were not assigned to a particular helper T cell subset using signature gene expression (Fig. 5a). This effect may be due to the limitations of scRNA-seq^31^, which are often difficult to distinguish from inherent heterogeneity in the CD4 T cell compartment^34^. We aimed to overcome these issues by using the single-cell ANnotation using Variational Inference (scANVI) deep generative model, which leverages high confidence seed labels based on signature marker genes to assign labels to unannotated cells^35^. We selected seed cells based on high expression of the gene signatures used in manual annotation (Fig. 5a, b), and also added lineage-specific transcription factor signature genes (*Tbx21* for Th1 and *Rorc* for Th17) to increase confidence in denoting these classical Th subsets. No cells of the nonclassical Th1-Th17 subset co-expressed *Tbx21* and *Rorc*, thus, scANVI seed labels for Th1-Th17 only required *Ifng* and *Il17a* expression. scANVI successfully annotated all previously unassigned cells and delineated similar clusters as manual annotation with signature genes (Fig. S6a), performing well at classifying cell types (Fig. S6b). Differential expression analysis of the scANVI-determined Th1 subsets showed similar top hits between samples (Fig. S6c) as manual annotation (Fig. S5e). scANVI classified many more cells as Th1-Th17 (Fig. S6d) compared to manual annotation (Fig. 6f), even reclassifying many Th17 as Th1-Th17. While Th1-Th17 cells were not readily detectable in naïve mice with manual annotation, scANVI revealed a small population of these cells in the mock vaccinated as well as a large cluster in the mucosal vaccinated sample (Fig. S6e). The pool of nonclassical CD4 T cells that do not fall into canonical *Tbx21*/*Ifng*-expressing Th1 or *Rorc*/*Il17a*-expressing Th17, therefore, is much larger than previously anticipated. These results show the promise and utility of recently developed tools in single cell transcriptomics and probabilistic models to study the heterogeneity of CD4 T cells in the context of vaccination and infection.

## Discussion

The recent success of the phase IIb clinical trial of the M72/AS01E protein subunit vaccine has raised the possibility that improved adjuvants and protein antigens may result in a protein subunit vaccine that confers complete protection against TB infection. In addition, protein subunit vaccines have potential for safe use in HIV-infected or otherwise immunocompromised individuals. Our results here provide mechanistic insight into a highly efficacious CDN-adjuvanted TB protein subunit vaccine. We found that intranasal vaccine-induced protection is evident 1 week post *M. tuberculosis* challenge in mouse lungs, correlates with a robust Th17 response, and is dependent on IL-17 for full efficacy. In contrast, BCG-induced protection is Th1-dominated and is delayed until the peak of the CD4 T cell response at 2-3 weeks post infection. While the STING-activated transcription factor STAT6 is dispensable for vaccine efficacy, type I IFN signaling is necessary for full protection elicited by our CDN vaccine. Although this vaccine elicits IFN-γ^+^ CD4 T cells when delivered through both the subcutaneous and intranasal routes, and IFN-γ plays an essential role in controlling *M. tuberculosis* infection, CDN vaccine efficacy requires a mixed T cell response requiring both IFN-γ and IL-17. Finally, single cell sequencing analysis of CD4 T cells at the peak of the T cell response reveal the emergence of a transcriptionally heterogeneous *Il17a*-expressing CD4 T cell cluster characterized by enhanced expression of functional markers is the most striking effect of mucosal vaccination. These results represent the first published single cell transcriptional analysis of naïve or vaccine-induced CD4 T cell responses during murine *M. tuberculosis* infection.

*M. tuberculosis* utilizes the ESX-1 secretion system to perforate host cell membranes to gain access to the cytosol and promote virulence, leading to dsDNA leak into the cytoplasm, activation of the cGAS/STING pathway, and subsequent induction of type I IFN and other pro-inflammatory cytokine responses^36^. BCG, however, lacks the ESX-1 system and is unable to permeabilize the phagosome^37^. Intriguingly, recombinant BCG vaccines that provide access to the cytosol provide superior protection^38^. Activation of cytosolic surveillance programs can therefore contribute to vaccine-induced immunity. To fully understand and improve upon the use of CDN adjuvants, it is essential to investigate the factors required for vaccine-induced protection downstream of STING activation. STING is primarily known for activating the transcription factor IRF3, resulting in type I IFN production, but transcription factors STAT6 and NF-κB can also be activated by STING^39^. We identified a partial role for type I IFN, but not STAT6, in mediating STING-dependent mucosal CDN vaccine efficacy. STING has also been shown to activate autophagy in a pathway independent of TBK1 and type I IFN induction^40,41^. It is therefore of great interest to further investigate the roles of type I IFN, NFkB and autophagy in eliciting CDN-adjuvanted mucosal vaccine protection.

While it is clear that IFN-γ from CD4 T cells is essential for host defense against M. tuberculosis, our understanding of IFN-γ-independent mechanisms of control has been limited. Although *Ifng*^−/−^ mice are extremely susceptible to *M. tuberculosis* infection in the presence or absence of the CDN-adjuvanted vaccine, both subcutaneous and intranasal administration induced protection that was only partially dependent on IFN-γ. Therefore, mucosal vaccination presents an exciting opportunity to explore CD4 memory T cell responses distinct from the classical Th1 response. While Th1 frequency has not proven as an effective correlate of protective immunity, boosting the mucosal memory response of a TB vaccine could synergize with IFN-γ-mediated immunity.

Our results contribute to a growing body of evidence that mucosal vaccines can elicit robust Th17 responses that augment Th1 responses induced by subcutaneous vaccination. In concordance with our findings, it was reported that an intranasal LT-IIb adjuvanted TB vaccine leads to a robust Th17 memory response in mice, and that IL-17 itself is required for mucosal vaccine efficacy^11^. However, the precise mechanism by which IL-17 specifically—or a Th17 T cell response more generally—protects against *M. tuberculosis* remains uncertain. We found that the efficacy of s.c. vaccination with either BCG or 5Ag/ML-RR-cGAMP is independent of IL-17. While it has been reported that subcutaneous administration of both a TB protein subunit vaccine and BCG results in IL-23-dependent polarization of inflammatory Th17 cells and protection^11,22^, this mechanism of control may be distinct from that of mucosal vaccination, as IL-23 may also be required for an IL-17-independent function during infection^42^. Further investigations are necessary to determine the role of IL-23 in CDN-adjuvanted vaccine efficacy.

One mechanism by which Th17 cells induce protection against TB may be the promotion of T cell localization to the lung parenchyma. While CXCR3 has primarily been described as a marker of protective Th1 parenchymal homing cells during *M. tuberculosis* infection^16^, its role in tissue-homing of pathogen-specific T cells in the context of vaccination has not been well-studied. Early and sustained lung parenchymal localization of protective CXCR3^+^ CD4 T cells may mediate our mucosal vaccine’s protection. As IL-17 is thought to mainly recruit neutrophils to the site of infection, it is of great interest to determine if and how this cytokine might induce and maintain this protective memory T cell response.

Although a robust Th1 response was seen in mock vaccinated mouse lungs at 3 weeks post infection, CD4 T cells in these mice expressed markers of terminal differentiation— KLRG1 and CX3CR1—and failed to infiltrate into lesions. Cytotoxic genes such as *Nkg7, Gzma*, and *Gzmb* were also expressed in a subset of these cells. Terminally differentiated Th1s are not associated with protection against TB^16,43^, and blood signature cytotoxic gene expression is anticorrelated with mouse susceptibility and progression from latent to active TB disease in humans^44^. Further investigation is necessary to determine if cytotoxic CD4s can mediate resistance to TB.

In contrast to the terminally differentiated Th1 marker expression seen in mock vaccinated mice, mucosal vaccination elicited higher expression of central memory T cell marker *Il7r* and activation/memory marker *Cd44* in the total CD4 and Th1 populations, and increased *Il1r1* expression specifically on IL-17-expressing T cells. Interestingly, i.n. vaccination also led to increased expression of *Tnfsf8*, which has been shown to be essential for IFN-γ-independent control of TB in a mouse model^15^ and implicated as a mediator of protection against human TB^33^. We thus find potential markers of both IFN-γ-dependent and IFN-γ-independent CD4 T cell memory function. These markers may also suggest mechanisms for protection: signaling through these molecules may contribute to enhanced activation, survival, function and memory development of CD4 T cells during infection. It will be crucial to evaluate whether these transcriptional changes result in memory responses that can provide enduring protection, past the point at which BCG-mediated immunity wanes.

While we had previously found that Th17 cells were induced by i.n. vaccination^18^, our latest findings reveal a surprisingly heterogeneous Th17 compartment, with 20-30% of *Il17a*-expressing cells in vaccinated mice also expressing *Ifng*. While CD4 T cell plasticity has been well documented in different contexts *in vivo*, it is uncertain whether this works to the host’s advantage. In fact, Th1-Th17 presence correlates with TB disease severity in humans^45^, and numerous studies have identified pathogenic IFN-γ^+^ Th17s as a driver of disease in mouse models of autoimmunity^13^. Furthermore, a previous study using adoptively transferred Th17s reported that the ability to coproduce IFN-γ limits Th17 protective capacity during *M. tuberculosis* infection in mice^12^. In our model, it is currently uncertain whether co-producing cells differ in protective capacity to classical Th1s or Th17s, and what cell types respond to IL-17 to mediate protection. Integrating single cell sequencing technology with spatial transcriptomics and cell surface proteomics provides exciting avenues for future investigations.

A recent mass cytometry study showed that the Th17 lineage maintains plasticity after *in vitro* differentiation^46^. Our scANVI analysis classified a surprisingly large proportion of both unassigned and *Il17a*-expressing “Th17” cells as Th1-Th17. This finding may indicate that Th1-Th17 cells are part of a larger pool of heterogeneous helper T cells, whose transcriptomes and effector functions may shift dynamically in response to inflammatory signals. Future fate-mapping studies could reveal whether CD4 T cell expression of IFN-γ and IL-17 changes over the course of vaccination, infection, and memory recall. While the plasticity in CD4 T cell phenotypes and function may be a hurdle in finding reliable correlates of vaccine correlates protection, true delineation of subtypes may be impossible, and it may be time to embrace heterogeneity in CD4 T cell-mediated protection for better TB vaccine design.

Our studies add to the growing body of literature proving that mucosal vaccination leads to the development of Th17 and Th1-Th17 T cell compartments. This heterogeneity distinguishes mucosal vaccination from the terminally differentiated Th1-dominated response elicited by systemic delivery of a vaccine. While intranasal delivery of the BCG vaccine also induces Th17 responses and protection, there remains a need to find a vaccine with an increased safety profile. Our vaccine has proven effective when administered twice as a booster to BCG and when delivered three times on its own^18^, inducing an IL-17-dependent response and protection that can outperform that of BCG. The studies presented here thus highlight the potentially profound impact of T cell diversity and lineage plasticity on antibacterial immune defense. Our findings suggest multiple avenues for improvement and optimism in designing improved vaccines against infectious respiratory diseases.

## Methods

### Ethics Statement

All procedures involving the use of mice were approved by the University of California, Berkeley Institutional Animal Care and Use Committee (protocol 2015-09-7979). All protocols conform to federal regulations, the National Research Council Guide for the Care and Use of Laboratory Animals, and the Public Health Service Policy on Humane Care and Use of Laboratory Animals.

### Vaccine Reagents

ML-RR-cGAMP was synthesized at Aduro Biotech as described previously^47^. AddaVax (InvivoGen, San Diego, CA) was used for the subcutaneous formulation of antigen and adjuvant as directed by the manufacturer. 5Ag fusion protein was provided by Aeras, H1 fusion protein was provided by Aeras and Statens Serum Institut, and peptide pools were provided by the NIH BEI Resources Repository.

### Mice

C57BL/6J (#000664), *Il17a*^*Cre*^ (#016879, referred to here as *Il17*^-/-^), *Ifng*^-/-^ (#002287) and *Stat6*^*-/-*^ (#005977) mice were obtained from The Jackson Laboratory (Bar Harbor, ME) and bred in-house. *Ifnar1*^-/-^ (*Ifnar*^-/-^, Jackson Laboratory strain #028288) mice were obtained from the Vance lab (University of California, Berkeley). Mice were sex- and age-matched for all experiments.

### Bacterial Culture

*M. tuberculosis* strain Erdman and *M. bovis* BCG (Pasteur) were grown in Middlebrook 7H9 liquid medium supplemented with 10% albumin-dextrose-saline (*M. tuberculosis*) or 10% oleic acid, albumin, dextrose, catalase (OADC) (BCG), 0.4% glycerol, and 0.05% Tween 80 or on solid 7H10 agar plates supplemented with 10% Middlebrook OADC (BD Biosciences) and 0.4% glycerol. Frozen stocks of *M. tuberculosis* and BCG were made from single cultures and used for all experiments.

### Vaccinations

5 μg ML-RR-cGAMP and 3 μg fusion antigen protein (5Ag or H1) were formulated in PBS for intranasal (i.n.) delivery or in 2% AddaVax in PBS for subcutaneous (s.c.) delivery. Mice were vaccinated three times at 4-week intervals with 20 μL i.n or with 100 μL s.c. at the base of the tail (50 μL on each flank). BCG-vaccinated mice were injected once with 2.5 x 10^5^ CFU/mouse in 100 μL of PBS plus 0.05% Tween 80 s.c. in the scruff of the neck.

### Challenge Experiments with *M. tuberculosis*

Twelve weeks after the initial vaccine injection, mice were infected via the aerosol route with *M. tuberculosis* strain Erdman. Aerosol infection was done using a nebulizer and full-body inhalation exposure system (Glas-Col, Terre Haute, IN). A total of 9 mL of culture diluted in sterile water was loaded into the nebulizer calibrated to deliver 50 to 100 bacteria per mouse as measured by CFU in the lungs 1 day following infection (data not shown).

### Pre-challenge ICS Assay

Heparinized blood lymphocytes were isolated 9 weeks post priming, processed as previously described^18^, stained with Live/Dead stain (Thermo Fisher Scientific, L34970), Fc receptor block, CXCR3, major histocompatibility complex (MHC) class II, KLRG1, and TCRγ/δ (BioLegend, 101319, 126522, 107606, 107606, and 118124, respectively), and CD4, CD8, CD90.2 and Ly6G (BD Biosciences, 564933, 563898, 561616 and 551460), and fixed/permeabilized with BD Cytofix/Cytoperm Fixation/Permeabilization Solution Kit (Thermo Fisher, 554714) before staining with IFN-γ (eBioscience, 12-73111-81) and IL-17 (BioLegend, 506904). Data were collected using a BD LSR Fortessa flow cytometer and analyzed using FlowJo Software (Tree Star, Ashland, OR).

### Post Challenge Cell Preparation

For bacterial enumeration, the largest lung lobe was homogenized in PBS plus 0.05% Tween 80, and serial dilutions were plated on 7H10 plates. CFU were counted 21 days after plating. The second largest lung lobe was fixed in 10% formalin for histological analysis or used for single cell RNA-sequencing. The remaining lung lobes were harvested into complete RPMI, dissociated, and strained through a 40 μm strainer. Cells were re-stimulated with no peptide or Ag85B peptide pools (2 μg/mL) with Protein Transport Inhibitor Cocktail (eBioscience, 00-4980-93) for 5 hours at 37C. Cells were washed and stained with antibodies used for pre-challenge ICS. Data were collected and analyzed as outlined above. Data presented are from Ag85B-stimulated samples for cytokine staining, and unstimulated samples for all other flow cytometric analyses.

### Immunofluorescence

Lung lobes were fixed in 10% formalin at room temperature for at least 24 hours. Histology was performed by HistoWiz Inc. (histowiz.com) using a Standard Operating Procedure and fully automated workflow. Samples were processed, embedded in paraffin, and sectioned at 4μm. Immunohistochemistry was performed on a Bond Rx autostainer (Leica Biosystems) with enzyme treatment using standard protocols. Rat monoclonal CD45R/B220 primary antibody (Biolegend, 103229) was used for immunohistochemistry, rat monoclonal CD3 primary antibody (Abcam) for immunofluorescence, and rabbit anti-rat secondary (Vector) DAPI was used to stain nuclei in Bond Polymer Refine Detection (Leica Biosystems) according to manufacturer’s protocol. After staining, sections were dehydrated and film coverslipped using a TissueTek-Prisma and Coverslipper (Sakura). Whole slide scanning (40x) was performed on an Aperio AT2 (Leica Biosystems). Slide images are available online at https://app.histowiz.com/shared_orders/6d2c7816-b292-42c4-9ee2-71ffe032957e/slides/. Hemotoxylin and eosin (H&E) and immunofluorescence staining was analyzed and scored using Olympus cellSens software by pathologists at the UC Davis Comparative Pathology Laboratory. Counts were generated using the QuPath software^48^. A single measurement classifier was generated for detection of cytoplasmic CD3 immunoreactivity within lesion. The classifier was applied to three equal-area ROI in each section of lung. For detection of cytoplasmic B220 immunoreactivity, classifiers were applied to the total sectional area of each lung section and the percentage of B220+ immune cells was determined. Lymphoid nodules were defined as discrete, nodular or ovoid, homogenous aggregates of lymphocytes with an area > 10000 um^2. Granulomas were defined as discrete, nodular aggregates of macrophages that efface pulmonary parenchyma, occasionally exhibit circumferential arrangement, and contain multinucleate giant cells.

### Single Cell RNA Sequencing

The second largest lung lobes were pooled from three mock or i.n. H1/ML-RR-cGAMP vaccinated mice, tissue dissociated in RPMI containing liberase and DNase I in gentleMACS C tubes using the gentleMACS dissociator and strained through a 70μm filter for sorting on a Sony SH-800 sorter using a 100μm nozzle. Sorted CD4 T cells were washed in RPMI and counted. 5000-6000 CD4 T cells per sample were used for single cell RNA sequencing (scRNA-seq) according to the 10x Genomics protocol. Briefly, single cell suspensions were partitioned into Gel Beads in emulsion (GEMs) using the 3’ Chromium Next GEM single Cell v3.1 system. 10x barcoded primers were used to reverse transcribe poly-adenylated mRNA to produce barcoded, full-length cDNA. Purified DNA was amplified, fragmented, and amplified again to attach Illumina adapter sequences by the Functional Genomics Laboratory at UC Berkeley. Libraries were sequenced on an Illumina NovaSeq SP 100SR and demultiplexed by the Vincent J. Coates Genomics Sequencing Laboratory at UC Berkeley. Reads were aligned to the mouse transcriptome mm10 with the 10x Cell Ranger analysis pipeline^49^, using the Savio computational cluster at UC Berkeley. After filtering, barcode counting, and UMI counting, the Scanpy toolkit^50^ was used to preprocess the data. The single cell variational inference (scVI)^32^ approach was utilized to project the data from both vaccinated and naïve mice into a shared lower-dimensional latent space. Uniform manifold approximation and projection (UMAP) and Leiden clustering were then applied on the *k-*Nearest-Neighbor graph of the latent space for visualization and clustering of the data. Leiden clusters that expressed high levels of myeloid, B cell, or CD8 T cell markers were excluded from the data before utilizing the scVI model a second time. The filtered data was annotated using signature genes from CD4 T cell subset in cellxgene (https://github.com/chanzuckerberg/cellxgene), a single-cell data exploration tool. Scanpy was used to visualize data and test differential expression of genes between samples and CD4 T cell subsets. The single-cell ANnotation using Variational Inference (scANVI)^35^ variant of scVI was used to assign annotations to unassigned cells based on high-confidence seed labels annotated on cellxgene. High-confidence seed cells were identified using a high threshold of expression as cutoffs. In the scANVI analysis, *Tbx21* and *Rorc* were also used to define classical Th1 and Th17, respectively.

### Statistical Analysis

Data are presented as mean values, and error bars represent SD. Symbols represent individual animals. The number of samples and statistical tests used are denoted in the legend of the corresponding figure for each experiment. Analysis of statistical significance was performed using GraphPad Prism 8 (GraphPad, La Jolla, CA), and p < 0.05 was considered significant.

## Acknowledgments

We thank Russell Vance for the kind gift of *Ifnar1*^*-/-*^ mice. We thank Kiran Magee and Lily McCann for assistance with mouse colony maintenance, Bianca Blackshire and Kiran Magee for assistance with *in vitro* experiments, and Dmitri Kotov for advice and assistance with scRNAseq experiments. We thank members of the Stanley and Cox labs for helpful discussions. We thank Aduro Biotech for the gift of cyclic-di-nucleotides. This work was supported by NIH grant T32 GM 7232-40 and NSF Graduate Research Fellowship DGE-1752814 (to E.V.D. and R.J.), and NIH 1R01AI113270-01A1 to SAS. This research used the Savio computational cluster resource provided by the Berkeley Research Computing program at the University of California, Berkeley (supported by the UC Berkeley Chancellor, Vice Chancellor for Research, and Chief Information Officer).

## Author contributions

SAS conceived of the study. SAS, RMJ, EVD and SM contributed to study design. RMJ, EVD, XN and AB performed experiments. GP performed analysis of histology. CX and NY assisted with scRNAseq analysis.

## Figure Legends

**Figure S1.**
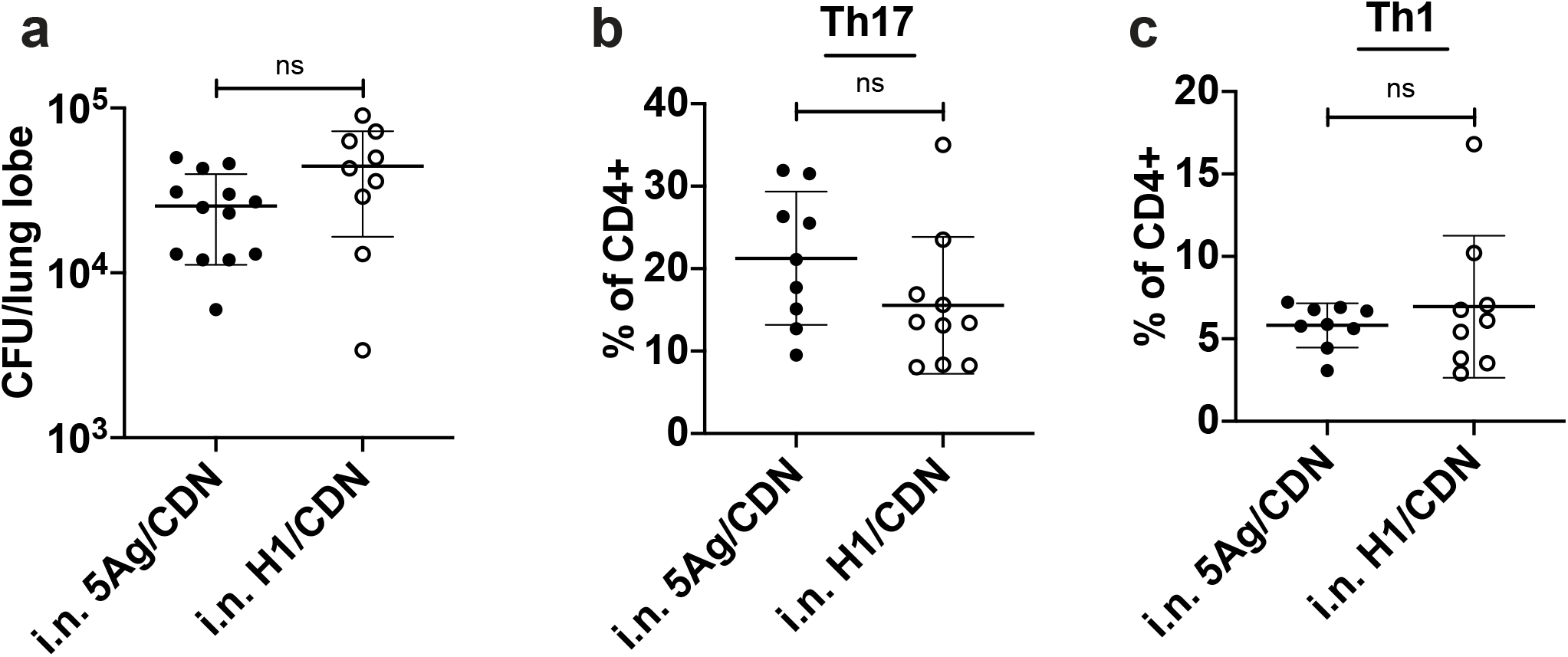
5Ag and H1 TB antigens provide equivalent efficacy in CDN-adjuvanted mucosal vaccines. Mice were administered three doses of intranasal 5Ag or H1 TB antigen adjuvanted with ML-RR-cGAMP. (a) At 4 weeks post challenge, bacterial burden was enumerated. (b) ICS for percentage of lung CD4 T cells that produce IL-17 (Th17), or (c) IFN-γ (Th1). Data are expressed as mean (± SD) of eight to ten mice per group from two independent experiments for each antigen. Mann-Whitney t test.

**Figure S2.**
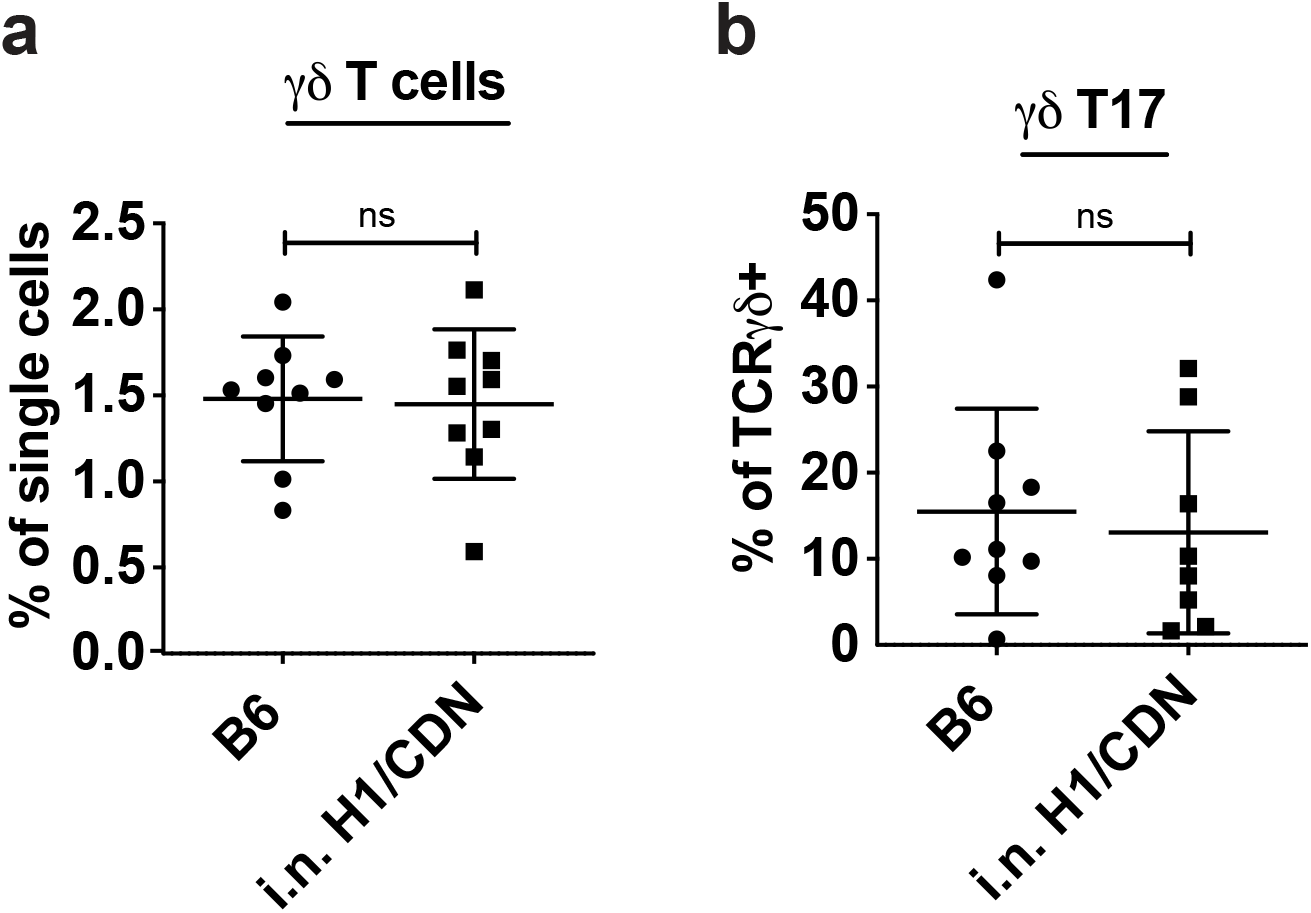
Intranasal immunization does not alter the frequency of TCRγ/δ or TCRγ/δ IL-17+ T cells. (a) Surface staining for percentage of live single cells that are TCRγ/δ+ in mock or i.n. H1/ML-RR-cGAMP vaccinated mouse lungs 4 weeks post M. tuberculosis challenge. (b) ICS for percentage of lung TCRγ/δ+ cells that produce IL-17 after restimulation ex vivo with recombinant Ag85b peptide pool. Data are expressed as mean (± SD) of eight to ten mice per group from two independent experiments; ICS data are representative of experiments with or without peptide restimulation. Mann-Whitney t test.

**Figure S3.**
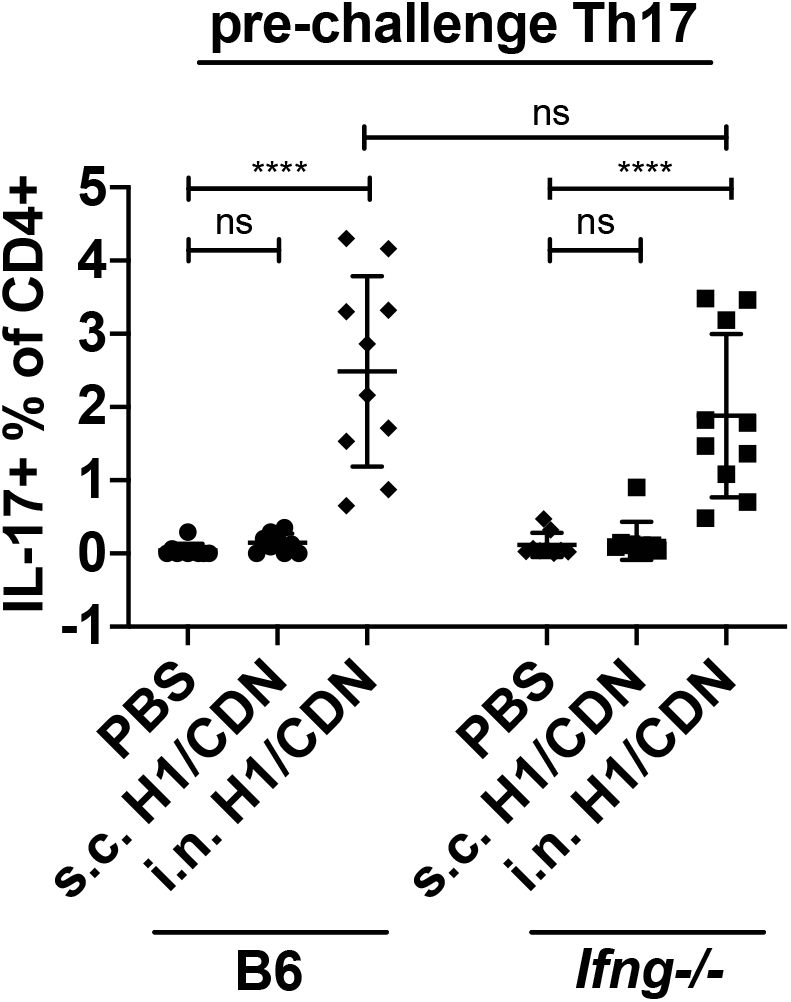
Intranasal vaccine-induced Th17 generation is unaffected by IFN-γ deficiency. Pre-challenge ICS of ex vivo restimulated PBMCs from blood of vaccinated B6 or Ifng-/-mice for Th17 percentage of CD4 T cells one week after the second vaccine boost. Data are expressed as mean (± SD) of eight to ten mice per group from two independent experiments. Mann-Whitney t test p values; ****p≤ 0.0001.

**Figure S4.**
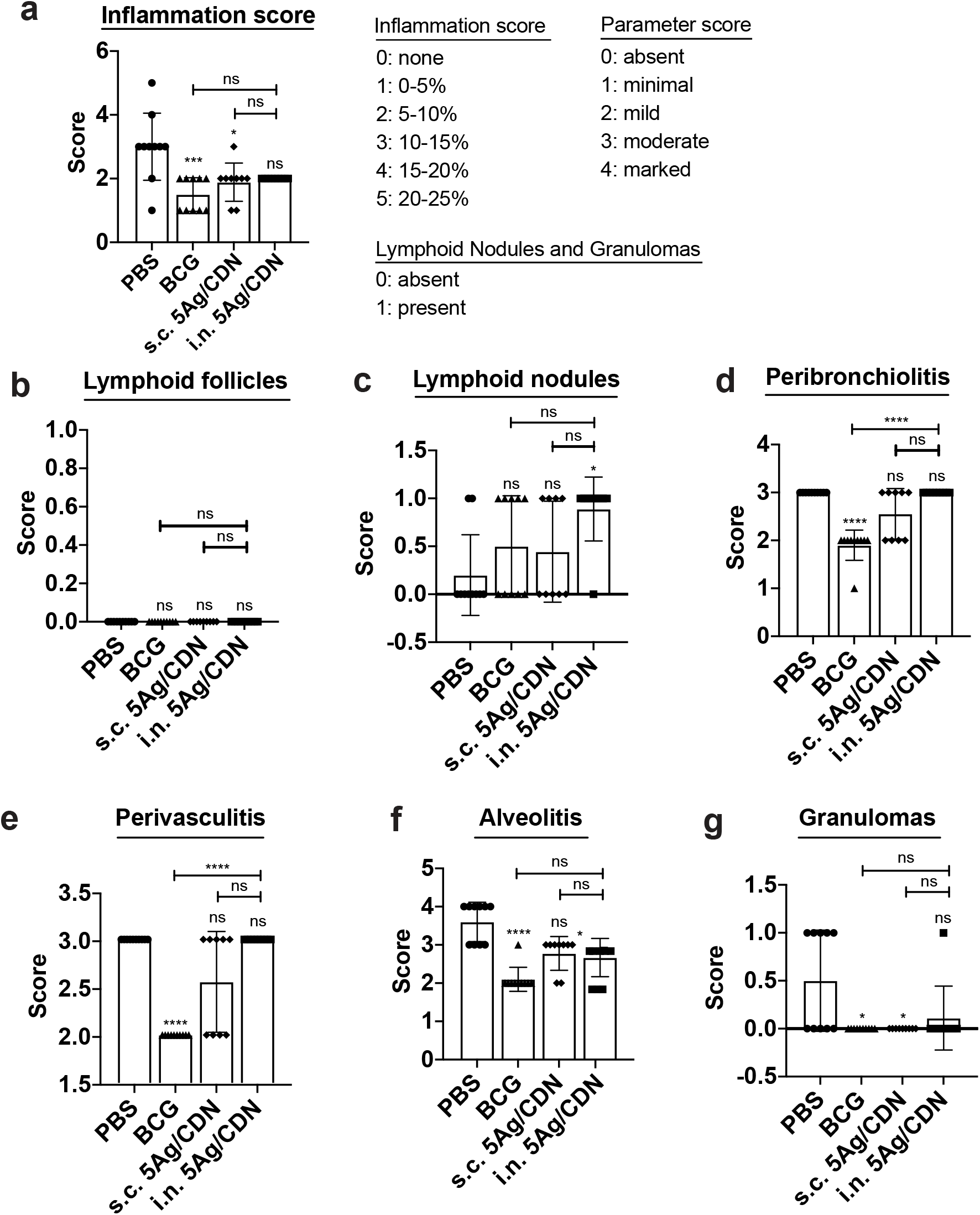
Semi-quantitative histopathological analysis of vaccinated mouse lungs. Formalin-fixed paraffin-embedded sections of mouse lungs 4 weeks post M. tuberculosis challenge were stained with H&E and assessed using a semi-quantitative scoring method for (a) inflammation area, (b, g) lymphoid follicle, lymphoid nodule, or granuloma presence, and (d-f) localized inflammation. Data are expressed as mean (± SD) of eight to ten mice per group from two independent experiments. Kruskal Wallis test followed by Dunn multiple comparison posthoc p-values; *p ≤ 0.05, ***p ≤ 0.001, ****p ≤ 0.0001.

**Figure S5.**
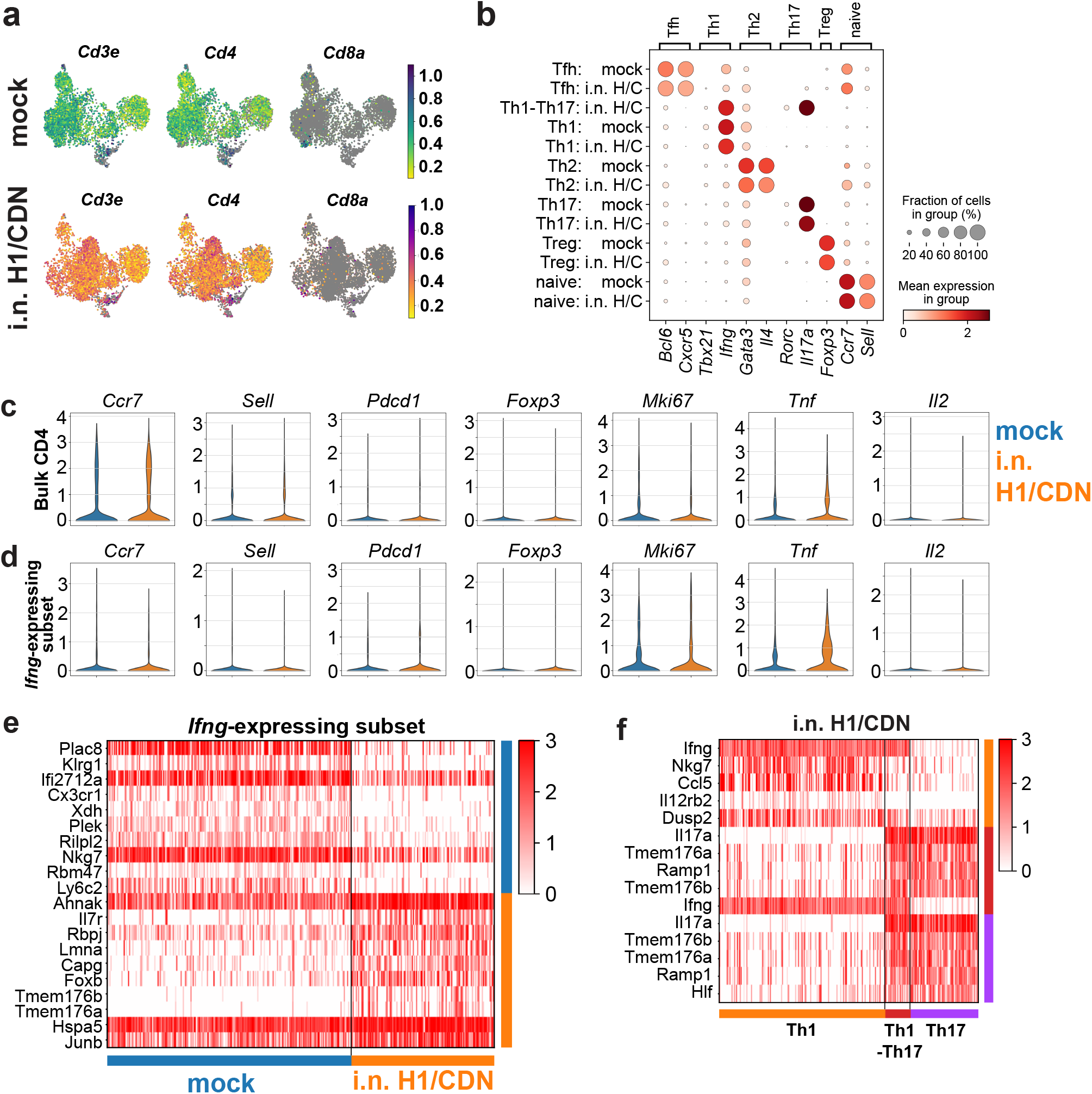
Single cell transcriptional analysis of mock and i.n. H1/ML-RR-cGAMP vaccinated mouse lung CD4 T cells. (a) UMAP plots of T cell marker expression in naïve (mock) or i.n. H1/ML-RR-cGAMP (i.n. H1/CDN) vaccinated mouse CD4 T cells. CD4 subtype signature gene expression in naïve or i.n. H1/ML-RR-cGAMP (i.n. H/C) vaccinated mice. (c) Gene expression in bulk and (d) Ifng-expressing CD4 T cell populations. (e) Top differentially expressed genes in Ifng+ CD4 T cells. (f) Top differentially expressed genes in Ifng+ Il17a-(Th1), Ifng+ Il17a+ (Th1-Th17), and Ifng-Il17a+ (Th17) subsets from i.n. vaccinated mouse lungs. T test with overestimated variance.

**Figure S6.**
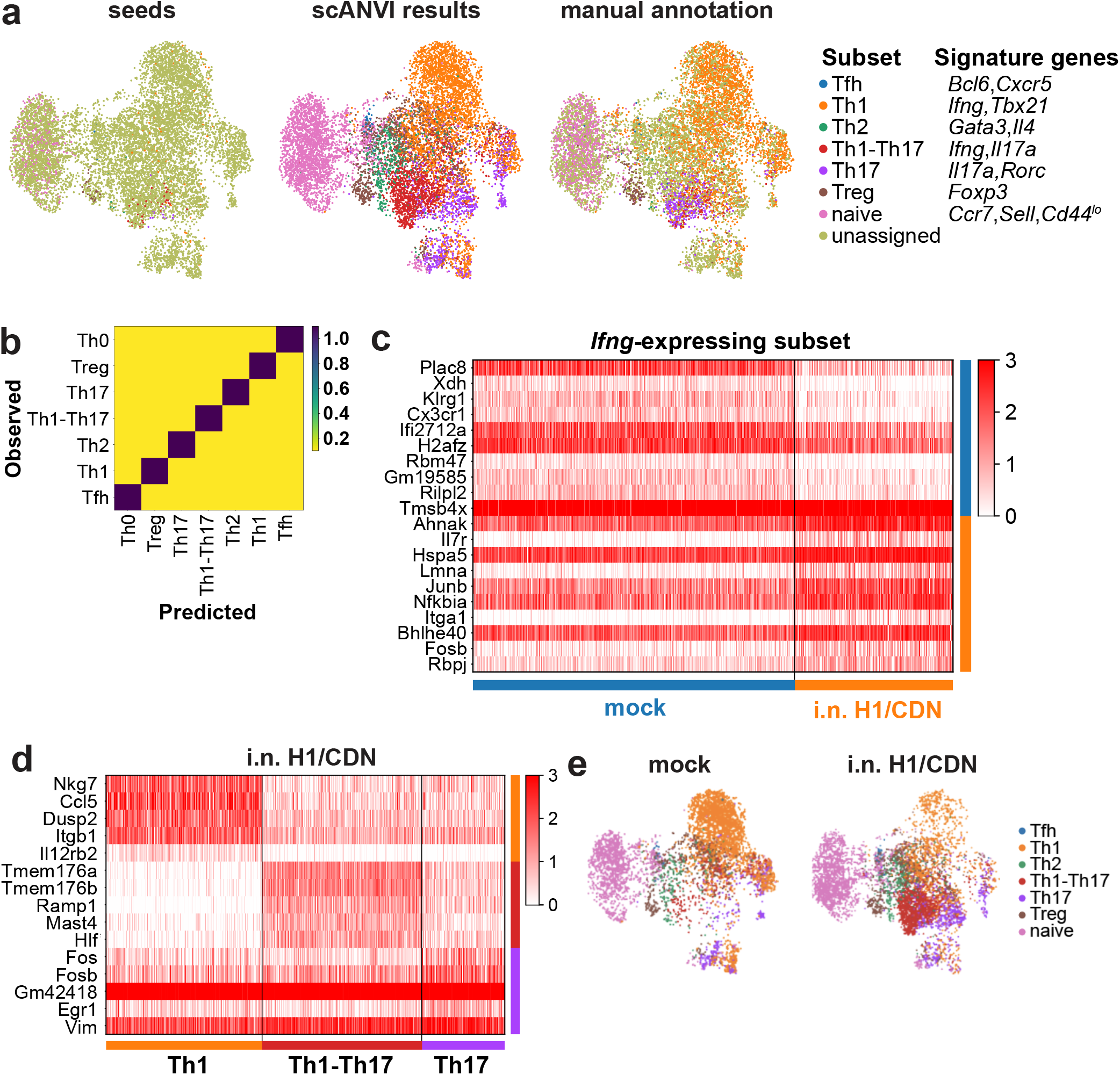
scANVI predicts T cell subset labels for unannotated cells. (a) UMAP plots of high confidence seed labels, scANVI annotations, and manual annotations used for analysis in Fig. 5 and 6 based on signature gene expression. (b) scANVI confusion matrix of observed and predicted labels to describe the model’s performance. (c) Top differentially expressed genes in scANVI-predicted Th1 cells in naïve (mock) or i.n. H1/ML-RR-cGAMP (i.n. H1/CDN) vaccinated mouse lung CD4 T cells. (d) Top differentially expressed genes in scANVI-predicted Th1, Th1-Th17, and Th17 subsets from i.n. vaccinated mouse lungs. (e) UMAP plots of scANVI-predicted CD4 T cell subsets from naïve or i.n. H1/ML-RR-cGAMP vaccinated mouse lungs. T test with overestimated variance.

